# Evidence for Involvement of *WDPCP* Gene in Alcohol Consumption, Lipid Metabolism, and Liver Cirrhosis

**DOI:** 10.1101/2023.04.11.536418

**Authors:** Felix O’Farrell, Benjamin Aleyakpo, Rima Mustafa, Xiyun Jiang, Rui Climaco Pinto, Paul Elliott, Ioanna Tzoulaki, Abbas Dehghan, Samantha H. Y. Loh, Jeff W. Barclay, L. Miguel Martins, Raha Pazoki

## Abstract

Alcohol consumption continues to cause a significant health burden globally. The advent of genome-wide association studies has unraveled many genetic loci associated with alcohol consumption. However, the biological effect of these loci and the pathways involved in alcohol consumption and its health consequences such as alcohol liver disease (ALD) remain to be elucidated. We combined human studies with model organisms *Drosophila melanogaster* and *Caenorhabditis elegans* to shed light on the molecular mechanisms underlying alcohol consumption and the health outcomes caused by alcohol intake. Using genetics and metabolite data within the Airwave study, a sample of police forces in the UK, we performed several analyses to identify changes in circulating metabolites that are triggered by alcohol consumption. We selected a set of genes annotated to genetic variants that are (1) known to be implicated in alcohol consumption, (2) are linked to liver function, and (3) are associated with expression (cis-eQTL) of their annotated genes. We used mutations and/or RNA interference (RNAi) to suppress the expression of these genes in *C. elegans* and *Drosophila*. We examined the effect of this suppression on ethanol consumption and on the sedative effects of ethanol. We also investigated the alcohol-induced changes in triacylglycerol (TGA) levels in *Drosophila* and tested differences in locomotion of *C. elegans* after acute exposure to ethanol. In human population, we found an enrichment of the alcohol-associated metabolites within the linoleic acid (LNA) and alpha linolenic acid (ALA) metabolism pathway. We further showed the effect of *ACTR1B* and *MAPT* on locomotion *in C. elegans* after exposure to ethanol. We demonstrated that three genes namely *WDPCP, TENM2* and *GPN1* modify TAG levels in *Drosophila*. Finally, we showed that gene expression of *WDPCP* in human population is linked to liver fibrosis and liver cirrhosis. Our results underline the impact of alcohol consumption on metabolism of lipids and pinpoints *WDPCP* as a gene with potential impact on fat accumulation upon exposure to ethanol suggesting a possible pathway to ALD.

## Introduction

Alcohol consumption is a major public health concern and is responsible for over 5% of the global burden of disease ^1^. It has been known for a long time that excessive drinking leads to a range of pathologies including alcohol liver disease (ALD). Some of the known biological changes involved in ALD are acceleration of hepatic lipogenesis in which *ADH* and *CYP2E1* genes are known to play a role ^2^.

Advances in genomics within the last two decades have resulted in a boost in our understanding of the mechanism of diseases through agnostic approaches such as genome-wide association studies (GWAS) that revealed numerous genetic loci linked to complex diseases. Recently, through GWAS we have identified genetic variants in the form of single nucleotide polymorphisms (SNPs) that are associated with alcohol consumption ^3^ as well as with circulating liver enzymes ^4^ in the European populations. Some of the identified genes (e.g. *ADH*, *KLB*, *DRD2*) have been investigated for better understanding of their involvement in alcohol consumption and health consequences such as ALD. However, the biological effect of most alcohol associated genes remain to be elucidated.

In this study, we aimed to shed light into the biological pathways involved in alcohol consumption and its effects towards ALD. We investigated metabolic changes in alcohol consumption and the pathways involved. We subsequently investigated biological effect of alcohol associated genes in ethanol exposed *C. elegans* and *Drosophila* to generate knowledge that could ultimately be used to develop new strategies for the prevention and treatment of ALD.

## Methods

### Population analysis

In the current study, we followed a multi-stage approach using population-based studies and model organisms to better understand pathways involved in alcohol consumption and its health consequences. We used data from the Airwave Health Monitoring Study ^5^, an occupational cohort of 53,116 police officers and staff ages 18 years and over across the UK. The Airwave Health Monitoring Study was approved by the National Health Service Multi-site Research Ethics Committee (MREC/13/NW/0588). Detailed information about the Airwave population, metabolic assays, data processing, metabolite annotation as well as genotyping and imputation is included in the supplementary methods.

We performed an agnostic association analysis between alcohol consumption and targeted metabolites and then investigated causality of the alcohol-metabolite associations and performed pathway analysis on the metabolites identified to have changed in the body due to alcohol consumption. We investigated known alcohol associated genes in *C. elegans* and *Drosophila* to pinpoint the genes involved in health consequences of alcohol consumption. Finally, we performed secondary analysis on the main results using bioinformatics approaches to provide a more comprehensive picture of the biological role of the genes involved in alcohol consumption. See below for a more detailed description of the above mentioned methods.

#### Analysis of the Metabolome

We analysed the association between alcohol consumption and circulating metabolites within the Airwave sample. Metabolomics features were obtained from data acquired by the National Phenome Centre (NPC) using liquid-chromatography / mass spectrometry), covering a wide range of hydrophilic and lipid metabolite classes. We excluded unannotated features from the analysis. To determine over-representation of alcohol-associated metabolites in known metabolic pathways, we performed pathway enrichment analysis using all annotated alcohol-associated metabolomics features. To this end, the Kyoto Encyclopedia of Genes and Genomes (KEGG) ^6^ and The Small Molecule Pathway Database (SMPDB) ^7^ were used. In addition, we performed association analyses between our alcohol-associated metabolites and known alcohol-associated SNPs and further performed causal inference analyses on these results using the Inverse variance weighted two-sample Mendelian randomization (MR) analysis. The aim of the MR analysis was to identify changes in circulating metabolites caused by alcohol consumption. MR is a causal inference method in observational studies that mimics randomised controlled trials (RCTs) by taking advantage of random assortment of alleles at conception. It uses instrumental variables (i.e. genotype status) that are robustly associated with an exposure of interest as a randomisation tool occurring naturally at conception ^8^.

#### Gene Selection for model organisms

We selected a list of 105 alcohol-related SNPs from recently conducted GWAS of alcohol consumption ^9, 10^. SNPs were selected if they presented a *P*-value lower than a GWAS significance threshold of 5 × 10^-8^ in their association with alcohol consumption. We sought for pleiotropic effect of these SNPs with liver function, a direct health burden of alcohol consumption, using our recently published GWAS of circulating liver enzymes ^4^. To account for multiple testing, a corrected *P*-value threshold of <0.00048 was used for the association with liver enzymes. This corresponds to a nominal *P*-value (0.05) that has been adjusted for the number of alcohol-related SNPs (n=105) using the Bonferroni method ^11^. Of the 105 SNPs associated with alcohol consumption, 43 SNPs were associated with at least one of the three liver enzymes alanine transaminase (ALT), alkaline phosphatase (ALP), and gamma-glutamyl transferase (GGT). We additionally used eQTL data within the GTEx database ^12^ to identify SNPs demonstrating effect on gene expression of their nearest genes (a cis-eQTL effect). We eventually selected 24 SNPs that showed evidence of a statistically significant eQTL effect (**Table 1**).

**Table 1.** Overview of the genetic variants with effect on alcohol consumption, liver enzymes and gene expression.

| Alcohol SNP * | Annotated Gene | eQTL tissue | eQTL effect size§ | eQTL P value† | Drosophila ortholog‡ | FlyBase score out of 15§ | Wormbase: C.elegans ortholog (score)‡ | Drosophila strains | C. elegans strain |
| --- | --- | --- | --- | --- | --- | --- | --- | --- | --- |
| rs823114 | NUCKS1 | Thyroid | -0.16 | $5.80 \times 10^{-18}$ | - | - | - | - | - |
| rs1260326 | GCKR | Thyroid | 0.22 | $1.80 \times 10^{-08}$ | - | - | - | - | - |
| rs2178197 | GPN1 | Brain-Cerebellar Hemisphere | -0.31 | $6.50 \times 10^{-08}$ | cg3704 | 13 | *gop-2* (9) ¶ | CG3704 (55294) | RG5036 *gop-2* (*gk5528*), |
| rs13032049 | WDPCP | Adipose-Subcutaneous | -0.16 | 0.0003 | *frtz* | 13 | - | Frz (55649) | - |
| rs11692435 | ACTR1B | Thyroid | 0.89 | $1.50 \times 10^{-80}$ | *arp1* | 12 | *arp-1* (8) ¶ | Lethal, not studied | FX4735 *arp-1* (*tm4735*) |
| rs12646808 | MSANTD1 | Thyroid | -0.25 | $1.30 \times 10^{-09}$ | cg18766 | 8 | - | - | - |
| rs11940694 | KLB | Liver | 0.22 | $4.00 \times 10^{-05}$ | cg9701 | 6 | *klo-1* (5), *klo-2* (5) | - | **Not studied** |
| rs1229984 | ADH1B | Esophagus-Gastroesophageal Junction | -1.4 | $1.90 \times 10^{-16}$ | *fdh* | 7 | *adh-5* (5) | - | **Not studied** |
| rs10078588 | TENM2 | Thyroid | 0.29 | $2.80 \times 10^{-15}$ | *ten-m* | 13 | *ten-1* (10) ¶ | TenM (29390) | VC518 *ten-1* (*ok641*), |
| rs34060476 | MLXIPL | Pancreas | 0.51 | $4.50 \times 10^{-18}$ | *mondo* | 12 | *mml-1* (9) | Mondo (27059) | RB954 *mml-1* (*ok849*) |
| rs10249167 | ARPC1B | Thyroid | 0.3 | $1.80 \times 10^{-25}$ | *arpc1* | 13 | *arx-3* (10) ¶ | Arpc1 (31246) | VC3166 *arx-3* (*ok1122*) |
| rs988748 | BDNF | Brain - Nucleus accumbens basal ganglia | 0.22 | $9.70 \times 10^{-06}$ | - | - | - | - | - |
| rs2071305 | MYBPC3 | Whole blood | -0.13 | $2.30 \times 10^{-11}$ | *hbs* | 1 | - | - | - |
| rs7121986 | DRD2 | Esophagus - Muscularis | 0.26 | $3.50 \times 10^{-08}$ | *dop2r* | 9 | *dop-2* (6), *dop-3* (6) | - | **Not studied** |
| rs10876188 | SLC4A8 | Cells-Cultured fibroblasts | -0.12 | $1.80 \times 10^{-07}$ | *ndae1* | 12 | *abts-1* (9) | Lethal, not studied | RB1381 *abts-1* (*ok1566*) |
| rs7958704 | SCN8A | Nerve - Tibial | -0.25 | $2.90 \times 10^{-10}$ | *para* | 12 | *cca-1* (1), *unc-77* (1), *egl-19* (4) | Para (33923) | JD21 *cca-1* (*ad1650*) |
| **rs12312693** | *STAT6* | Brain - Frontal Cortex BA9 | 0.23 | $8.00 \times 10^{-05}$ | *stat92e* | 9 | *sta-2* (4), *sta-1* (7) | - | RB796 *sta-1* (*ok587*) |
| **rs11625650** | *KIF26A* | Whole blood | -0.26 | $9.10 \times 10^{-12}$ | *cg14535* | 7 | *vab-8* (2) | - | NG2484 *vab-8* (*gm84*) |
| **rs1421085** | *FTO* | Muscle - Skeletal | 0.14 | $7.50 \times 10^{-08}$ | - | - | - | - | - |
| **rs11648570** | *PMFBP1* | Esophagus - Mucosa | -0.29 | $9.00 \times 10^{-05}$ | *cg12702* | 1 | - | - | - |
| **rs3803800** | *TNFSF13* | Brain - Cortex | -0.2 | $2.50 \times 10^{-08}$ | *egr* | 3 | - | - | - |
| **rs1053651** | *TCAP* | Muscle - Skeletal | -0.11 | $4.90 \times 10^{-11}$ | - | - | - | - | - |
| **rs1991556** | *MAPT* | Brain - Cerebellum | -0.28 | $1.80 \times 10^{-06}$ | *tau* | 7 | *ptl-1* (8) | - | RB809 *ptl-1* (*ok621*) |
| **rs4815364** | *ACSS1* | Brain - Cerebellar Hemisphere | -0.47 | $1.70 \times 10^{-12}$ | *accoas* | 6 | - | - | - |

\* Alcohol SNPs associated with liver enzymes at replication P value <0.000476. Liver enzyme GWAS summary statistics obtained from Pazoki et al 2021.
† Normalised effect size and P-value obtained from GTEx portal. ‡ Orthologs were extracted from FlyBase and WormBase. § number of FlyBase tools that support the gene-pair relationship out of a total number of tools (n=15) that computed relationships between *Homo sapiens* and *Drosophila melanogaster*.
¶ Genes for which mutations were homozygous lethal and as such, heterozygote mutations balanced by chromosomal translocations were instead analysed.
eQTL: expression quantitative trait loci.

#### Ortholog Selection

The 24 SNPs with cis-eQTL effect with their annotated genes were matched to their orthologs in Drosophila using FlyBase (DIOPT online tool version 8.5/9.0; beta; http://www.flyrnai.org/diopt). To ensure selection of the most credible orthologs, we used scores calculated in FlyBase. This database provides a number of approaches that support the gene-pair relationship out of a total number of tools that computed relationships between *Homo sapiens* and *Drosophila*. Genes with a score of >12 were shortlisted for further analysis in *Drosophila*. Nineteen *Drosophila* orthologs were identified of which eight had a score ≥ 12 including *ARPC1B* (*arpc1*), *ACTR1B (arp1), GPN1* (*CG3704*), *WDPCP* (*frz*), *MLXIPL* (*mondo*), *SLC4A8 (ndae1), SCN8A* (*para*), and *TENM2* (*Ten-m*).

To identify potential worm orthologs, the SNPs with cis-eQTL effect with their annotated genes were sought for their worm orthologs within WormBase (https://wormbase.org/). BLASTp analysis results of human protein sequence against *C. elegans* protein database were obtained in WormBase version WS280 using data from *C. elegans* Sequencing Consortium genome project (PRJNA13758). The human protein sequence for the protein encoded by the genes under this study was extracted from UniProt (https://www.uniprot.org/). Worm genes with best alignment with the human protein sequence indicated by an Expect value (E value) < 1 × 10^-5^ and the highest BLASTp score (bits) were moved forward. The E value represents the number of alignments that could be found in similarity to the protein sequence by chance. Using WormBase, 13 worm orthologs were identified including *ACTR1B* (*arp-*1), *ARPC1B* (*arx-3*), *GPN1* (*gop-2*), *MLXIPL* (*mml-1*), *STAT6* (*sta-1*), *TENM2* (*ten-1*), *KIF26A* (*vab-8*), *SLC4A8* (*abts-1*), *MAPT* (*ptl-1*) and *SCN8A* (cca-1). We excluded *KLB* (*klo-1*), *DRD2* (*dop-2*), and *ADH1B* (*adh-5*) from the *C. elegans* experiments as similar studies investigating these genes already exist ^13–15^.

We did not identify any fly or worm orthologs for *NUCKS1*, *GCKR*, *BDNF*, *FTO*, *TCAP*. Additionally, no worm ortholog was found for *WDPCP, MSANTD1, MYBPC3, PMFBP1, TNFSF13,* and *ACSS1*.

### Drosophila

#### Genetics and Drosophila melanogaster strains

All *Drosophila* stocks and crosses were maintained on standard cornmeal agar media at 25°C on 12/12 hour light/dark cycles. The following strains were used as positive control: *w^berlin^; hppy^17–51^; + (hppy* mutant) and *w^berlin^; hppy^17–51^; hppy III* (*hppy* mutant with a rescue genomic construct) (gifts from Prof. Ulrike Heberlein, Janelia Research Campus, Virginia, USA), RNAi lines were obtained from Bloomington *Drosophila* Stock Center. All lines used were backcrossed to *w^1118^* or [*v*]*^w1118^* (RNAi lines). All RNAi expressions were driven by *da*Gal4. All the experiments on adult flies were performed using males.

#### Drosophila Ethanol Consumption Assay

The CApillary FEeder (CAFE) assay ^16^ was used to measure Ethanol consumption. Eight male flies were placed into an experimental vial (8 cm height, 3.3 cm diameter) containing 6 microcapillary tubes (BRAND® disposable BLAUBRAND® micropipettes, intraMark, BR708707, with 1 µL marks), each containing 5 µL of liquid food. Liquid food was prepared by dissolving 50 mg of yeast granules in 1 ml of boiling water by vortexing, followed by brief centrifugation. Then, 40 mg of sucrose (Sigma‒ Aldrich, 84097) was added to 800 µL of the dissolved yeast mixture, followed by vortexing. The microcapillary tubes were filled with liquid food up to the 5 µL mark. Ethanol food consists of normal food supplemented with 15% Ethanol. Each experiment consisted of 5 experimental vials per genotype with each vial containing normal food (3 capillaries) and Ethanol food (3 capillaries). The flies were acclimatised in the experimental vial without any food for 2 hours prior to the start of the experiment. This step was also used to incentivise the flies to eat once the food was introduced. The experimental vials were placed in a plastic box with a cover to control humidity. The flies were allowed to feed for 19 hours, after which the amount consumed (in mm) was measured with a digital calliper (Dasqua Bluetooth Digital Calliper 12″/300 mm, 24108120). The total amount of food consumed was calculated using the formula:

Food uptake (µl) = (Σ measured distances between 3 microcapillary tubes (mm))/14.42 mm (1 µl in measured distance) ⁄ 8 flies.

#### Drosophila Ethanol sedation Assay

Fly sedation assay was performed as previously described ^17^. Briefly, 8 flies were transferred to a 25mm x 95mm transparent plastic vial in between two cotton plugs. A piece of cotton plug at the base of the vial served as a stable surface to observe the flies and another plug was used to cap the vial and deliver the ethanol. 500ul of 100% Ethanol was added to the side of the cotton plug facing the flies. Sedation was observed manually as ST50, which is the time in minutes it takes for 50% of the flies in a sample vial to become sedated. Sedation events are recorded when the flies become inactive and lay on their backs for over 10 seconds.

#### Drosophila TAG measurements

Eight male flies of the indicated genotypes were placed into an experimental vial as described in ethanol consumption assay, with all 6 microcapillary tubes filled with normal food (5% sucrose + 5% yeast) or ethanol food (normal food + 15% ethanol), for 2 days. TAGs were assessed through colorimetric assays using 96-well microtiter plates and an Infinite M200Pro multifunction reader (TECAN). The assays were performed as previously described ^18^. Briefly, flies were homogenised in 110 µl of PBS + 0.05% Tween 20 (PBST) for 2 min on ice and immediately incubated at 70 °C for 10 min to inactivate endogenous enzymatic activity. A 35 µL fly homogenate sample and a glycerol standard (Sigma, no. G7793) were incubated together with either 35 µL of PBST (for free glycerol measurements) or 35 µL of TAG reagent (Sigma, no. T2449, for TAG measurements) at 37 °C for 60 min. After 3 min of centrifugation at full speed, 30 µL of each sample was transferred into a clear-bottom plate (two technical replicates per biological sample) together with 100 µL of free glycerol reagent (Sigma‒Aldrich, F6428) and incubated at 37°C for 5 min. TAG absorbance was divided by the protein concentration of the respective sample, which was measured by Bradford assay (Sigma‒Aldrich, B6916).

### C. elegans

#### Nematode strains and culture

All *C. elegans* strains were cultured on nematode growth medium (NGM) agar plates at 20°C using *Escherichia coli* OP50 as a food source. For the wildtype worms Bristol N2 strain was used. Genes for which mutations were homozygous lethal, heterozygote mutations balanced by chromosomal translocations were instead analysed.

#### Nematode RNA interference (RNAi)

RNAi experiments were performed on the NL2099 *rrf-3 (pk1426)* strain as previous described ^13, 19^. RNAi was achieved by feeding ^20^ using the ORFeome based RNAi library ^21^. In brief, HT115 RNAi bacterial clones were initially cultured in LB media with 100 µg/ml ampicillin and subsequently spotted in three 50 µl drops on 60 mm diameter NGM plates containing 1 mM isopropyl β-1-thiogalactopyranoside (IPTG) and 25 µg/ml carbenicillin. Plates were left to dry for 4-7 days before seeding to improve RNAi efficiency. Following seeding, five L3-L4 worms were added to each RNAi plate and cultured at 20°C until the F1 generation reached adulthood. Ethanol experiments were performed and analysed as described above and compared to worms fed with an empty RNAi feeding vector.

#### Nematode Behavioural assays

All ethanol experiments were performed at 20°C in a temperature-controlled room as previously described ^13, 19^. Behavioural assays were conducted on young adult hermaphrodites selected from sparsely populated NGM plates. Nematodes with *loss-of-function* mutations in worm orthologues of *ACTR1B* (*arp-*1), *ARPC1B* (*arx-3*), *GPN1* (*gop-2*), *MLXIPL* (*mml-1*), *STAT6* (*sta-1*), *TENM2* (*ten-1*), *KIF26A* (*vab-8*), *SLC4A8* (*abts-1*), *MAPT* (*ptl-1*) and *SCN8A* (cca-1) were acutely exposed to ethanol and the resultant effect on rate of locomotion (thrashes per minute) was quantified in Dent’s solution (140 mM NaCl, 6 mM KCl, 1 mM CaCl2, 1 mM MgCl2, 5 mM HEPES, pH 7.4 with bovine serum albumin at 0.1 mg/ml) by measuring thrashes per minute (one thrash defined as one complete movement from maximum to minimum amplitude and back) following 10 minutes exposure to the drug. Ethanol was mixed with Dent’s solution at a concentration of 400 mM, which has previously been shown to produce a ∼70% reduction in locomotion rate in wild-type worms ^19, 22^.

### Bioinformatics

#### Secondary analysis

To gain a better insight into the biological pathways involved in the link between alcohol consumption and liver damage, we used the genetic variants within the genes highlighted by our model organism experiments and performed a series of secondary analysis using human data. We explored publicly available data from the UK Biobank deposited in the Edinburgh Gene Atlas ^23^ using which we sought Phewas (Phenome-wide association analysis) databases to obtain association results between the genetic variants and 778 traits. We additionally used the genes highlighted by our model organism experiments to assess the causal effect of gene expression on liver conditions. Within these genes, the SNPs that have been identified to have cis-eQTL effect within the previously published studies were selected and used as MR instrument against liver conditions within the *twosampleMR* package in R.

#### Statistical analyses

Within the Airwave study sample, we performed a linear regression to study the association of alcohol consumption with each of the metabolomic features (Metabolome-wide association study; MWAS). We adjusted the statistical analysis for age, sex, smoking status, and salary class. To account for multiple testing and the high degree of correlation in metabolomics datasets, we used a permutation-based method to estimate the significance level of the associations ^24, 25^. For each metabolomics platform, a *P*-value threshold equivalent to adjusting to a 5% Family-Wise Error Rate (i.e., Bonferroni method) was computed. A series of hypergeometric tests implemented in the R package MetaboAnalystR ^26^ was used for pathway enrichment analysis where an FDR threshold of 0.05 was used as a significance threshold. To obtain an estimation for the association of known alcohol SNPs with our alcohol-associated metabolites to be used in the MR analysis, we performed linear regression analysis within the Airwave sample (see supplementary methods for details of the GWAS on metabolomics). Linear analyses between SNPs and metabolites were conducted for each metabolomic feature with adjustment for age, sex, and genetic principal components within a subsample of Airwave that included participants with both genetic and metabolite data (N=1,970). In *C. elegans,* locomotion rate was presented normalised as a percentage of mean thrashing rate of untreated worms measured each day. All worm data were expressed as mean ± S.E. with an N=30 individual worms. Locomotion rate significance was assessed by one-way analysis of variance (ANOVA) with post-hoc Tukey test for multiple comparisons. Statistical analyses of the *Drosophila* experimental data were performed using GraphPad Prism (www.graphpad.com). *Drosophila* data were presented as the mean values, and the error bars indicate ± SD.

## Results

To understand biological effects of alcohol consumption in human population, we investigated the circulating metabolites within the Airwave study sample using an agnostic approach which revealed association of 376 metabolites with alcohol consumption and alcohol-associated genetic variants (**Figure 1**). Pathway analysis on the identified metabolites showed that the linoleic acid (LNA) and alpha linolenic acid (ALA) metabolism pathway (LNA/ALA) within the Small Molecule Pathway Database (SMPDB) was enriched with alcohol-associated features (Pfdr=5.67 × 10^-3^). These features included Tetracosapentaenoic acid (24:5n-3; β=0.01; 95%CI=0.008,0.012; *P*-value=4.2 ×10^-12^), Eicosapentaenoic acid (β=0.009; 95%CI=0.007,0.011; *P*-value=2.4 × 10^-12^), Stearidonic acid (β=0.008; 95%CI=0.006,0.01; *P*-value=2.2 × 10^-9^), Arachidonic acid (β=0.007; 95%CI=0.005,0.009; *P*-value=9.7 × 10^-8^) and Adrenic acid (β=0.01; 95%CI=0.008,0.012; *P*-value=9.7 × 10^-8^). To assess the causality of these associations, we used alcohol-associated genetic variants against circulating metabolites within the Airwave sample using MR analysis (**Table 2**). This revealed possible causal association of alcohol consumption on changing circulating level for several lipid metabolites (TAGs, Diradylglycerols, Glycerophosphocholines, Sphingolipids), and an alkaloid (piperine). The most statistically significant causal association was observed with a triacylglycerol TG 60:2 (β=1.24; 95% CI=0.52,1.95; *P*-value=0.002).

**Figure 1:**
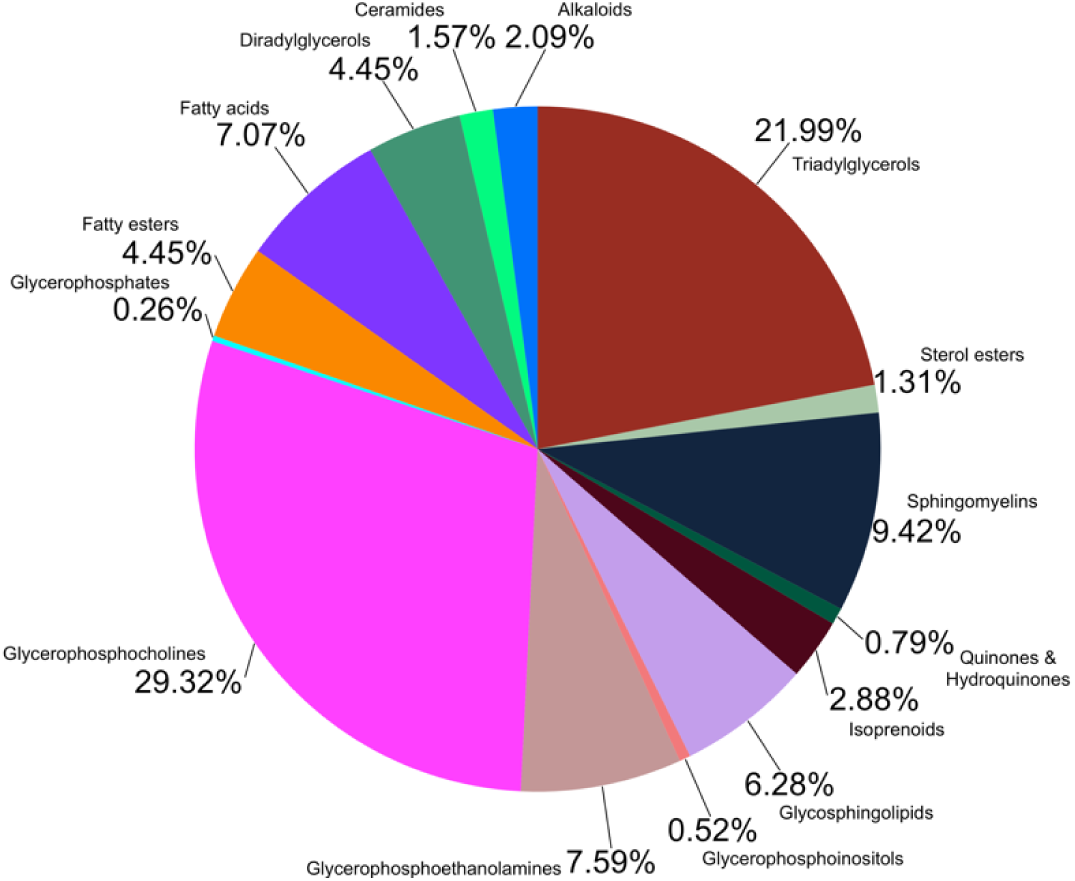
Association of alcohol consumption with circulating metabolites in the Airwave study. Pie chart illustrates percentages of metabolite classes (RefMet “Super class”) present that are significantly associated with alcohol consumption.

**Table 2.** Overview of the causal effect of alcohol on circulating metabolites using the inverse variance weighted two-sample Mendelian randomisation multiple instrument method.

| Metabolite | Name | Main class | Beta (95% CI) | P-value |
| --- | --- | --- | --- | --- |
| SLPOS_457.3339_0.7190 | CAR DC18:1 | Fatty esters | -1.13 (-2.14, -0.11) | 0.036 |
| SLPOS_579.5352_8.2395 | DG 34:0 | Diradylglycerols | 0.91 (0.05,1.77) | 0.047 |
| SLPOS_606.5548_8.3326 | DG 36:1 | Diradylglycerols | 0.92 (0.07,1.77) | 0.041 |
| SLPOS_745.5603_5.7424 | PC 33:2 | Glycerophosphocholines | 0.78 (0.07,1.5) | 0.040 |
| SLPOS_629.5424_6.9217 | PE 38:4 | Glycerophosphoethanolamines | 1.2 (0.15,2.25) | 0.032 |
| SHPOS_625.5174_2.8914 | PE 38:5 | Glycerophosphoethanolamines | 1.1 (0.1,2.1) | 0.039 |
| SLPOS_201.0516_0.6580 | Piperine | Alkaloids | 1.16 (0.1,2.22) | 0.040 |
| SHPOS_807.6348_4.4974 | SM 40:2;O2 | Sphingomyelins | 1.06 (0.15,1.96) | 0.028 |
| SHPOS_827.7093_0.5837 | TG 48:1 | Triradylglycerols | 0.95 (0.13,1.77) | 0.030 |
| SLPOS_898.7965_11.2257 | TG 53:1 | Triradylglycerols | 0.93 (0.1,1.75) | 0.036 |
| SLPOS_888.8079_10.7574 | TG 53:3 | Triradylglycerols | 0.96 (0.06,1.85) | 0.045 |
| SLPOS_607.5599_11.1277 | TG 54:2 | Triradylglycerols | 0.95 (0.19,1.71) | 0.019 |
| SLPOS_1027.7392_10.9051 | TG 54:3 | Triradylglycerols | 1.06 (0.21,1.91) | 0.020 |
| SLPOS_984.8508_11.7812 | TG 58:1 | Triradylglycerols | 1.12 (0.44,1.81) | 0.003 |
| SLPOS_687.6307_11.5996 | TG 58:2 | Triradylglycerols | 1.08 (0.27,1.9) | 0.014 |
| SLPOS_689.6461_11.7895 | TG 60:2 | Triradylglycerols | 1.24 (0.52,1.95) | 0.002 |

CI, Confidence Interval. Effect estimates and 95% CI is given for the inverse variance weighted method of Mendelian randomisation.

We used *Drosophila hppy* mutant, described to have an increased resistance to ethanol sedation ^27^, as positive control in all fly experiments. *Drosophila* with knock-down of *ARPC1B* (*arpc1*), *GPN1* (*CG3704*), *WDPCP* (*frz*), *MLXIPL* (*mondo*), *SCN8A* (*para*), and *TENM2* (*Ten-m*) together with *hppy* mutant were exposed to food supplemented with 15% Ethanol in a CAFE assay. A significant difference (**Figures 2A & 2B**) was observed between ethanol consumed (µL) by flies with RNAi knockdown of *TENM2* (*Ten-m*). Following exposure to ethanol vapour, the effect on Sedation Time 50% (ST50; minute) was quantified and we observed that in comparison to control (*hppy*), RNAi knockdown of fly orthologues of *WDPCP* (*frz*) showed a faster rate of sedation whilst *TENM2* (*Ten-m*), *GPN1* (*CG3704*), *ARPC1B* (*arpc1*) and *SCN8A* (*para*) showed a slower rate of sedation indicated by a higher ST50. *WDPCP* (*frz*) knockdown flies showed reduced TAG levels after exposure to ethanol food whilst *TENM2* (*Ten-m*) and *GPN1* (*CG3704*) knockdown flies showed a reduction in TAG levels with normal food (**Figure 2C**).

*C. elegans loss-of-function* mutants of *TENM2* (*ten-1)*, *KIF26A (vab-8)*, *SLC4A8 (abts-1)* and *SCN8A (cca-1)* showed significant differences in basal locomotion rate (**Supplementary Figure S1A).** After acute exposure to ethanol in *C. elegance loss-of-function* mutants, significant differences were identified in normalised locomotion rate of *ACTR1B* (*arp-*1), *ARPC1B* (*arx-3*) and *MAPT* (*ptl-1*) in comparison with Bristol N2 wild-type worms (**Figure 3A**). RNAi knockdown of these genes confirmed that in comparison to controls, RNAi knockdown of worm orthologues of *ACTR1B* (*arp-*1) and *MAPT* (*ptl-1*) and *ARPC1B* (*arx-3*) did not have any effect on basal locomotion rate (**Supplementary Figure S1B)** but RNAi knockdown of *ACTR1B* (*arp-*1) and *MAPT* (*ptl-1*) phenocopied the *loss-of-function* mutations after exposure to ethanol (**Figure 3B**).

**Figure 2:**
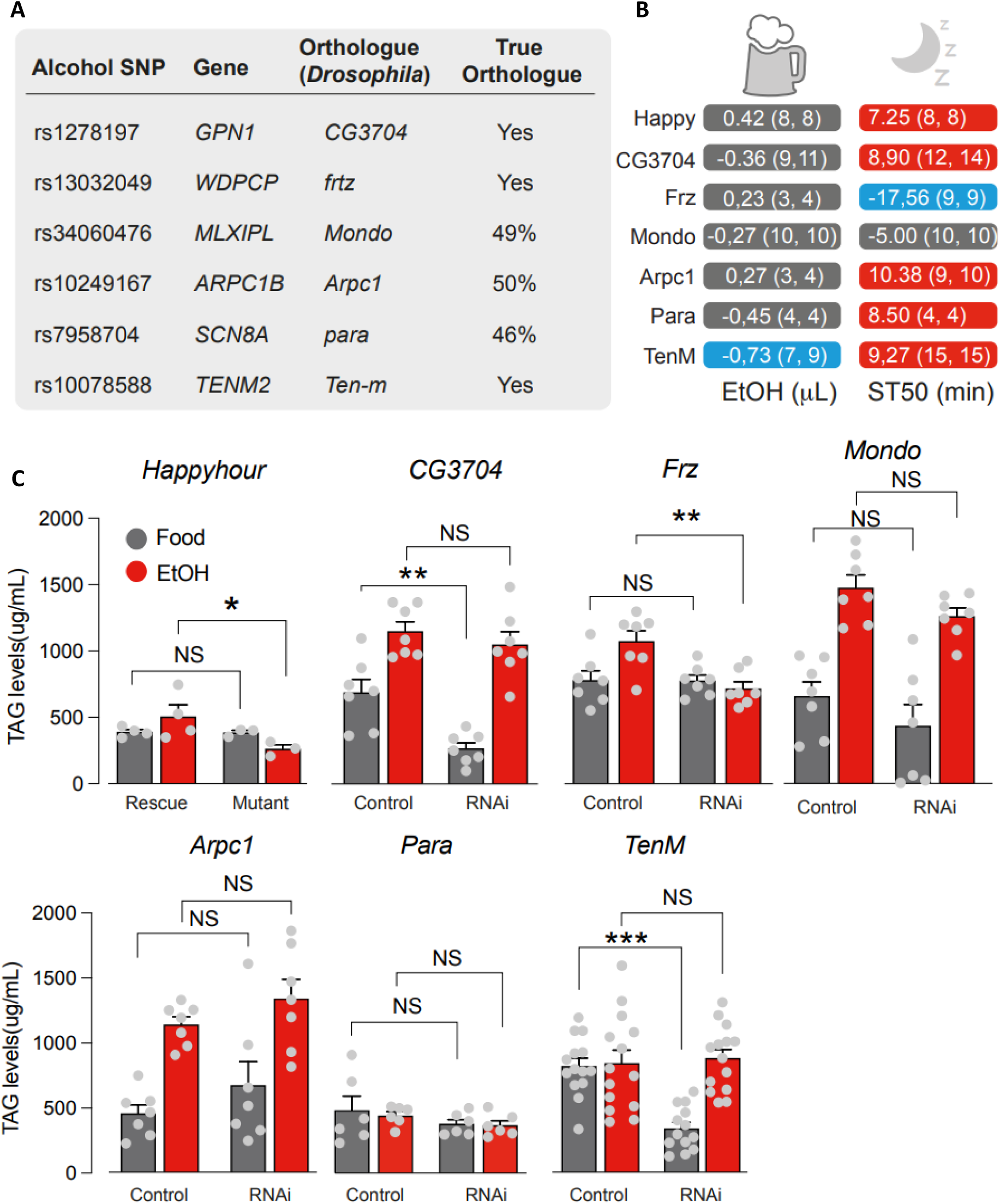
Analysis of alcohol intake and sedation in adult flies. (A) Mapping *Drosophila* orthologues of human genes involved in ethanol consumption. (B) Analysis of ethanol intake (left column) and sedation (right column) in the Drosophila RNAi lines. The numerical values show the difference between means (RNAi - Control). Values in parenthesis are the number of biological replicates for respectively, the control and the RNAi line. (C) Analysis of TAG levels in adult flies fed either with normal or ethanol-containing food (mean ± standard error of mean; asterisks, 2-way ANOVA with Tukey’s multiple comparisons test). The number of biological replicates per experimental variable (n) is indicated in either the respective figure or figure legend. No sample was excluded from the analysis unless otherwise stated. Blinding was not performed. Normality was assessed before deciding on which parametric or non-parametric test to use for inferential statistics. Statistical significance is indicated as * for P < 0.05, ** for P < 0.01, *** for P < 0.001, **** for P < 0.0001 and NS for P ≥ 0.05.

**Figure 3:**
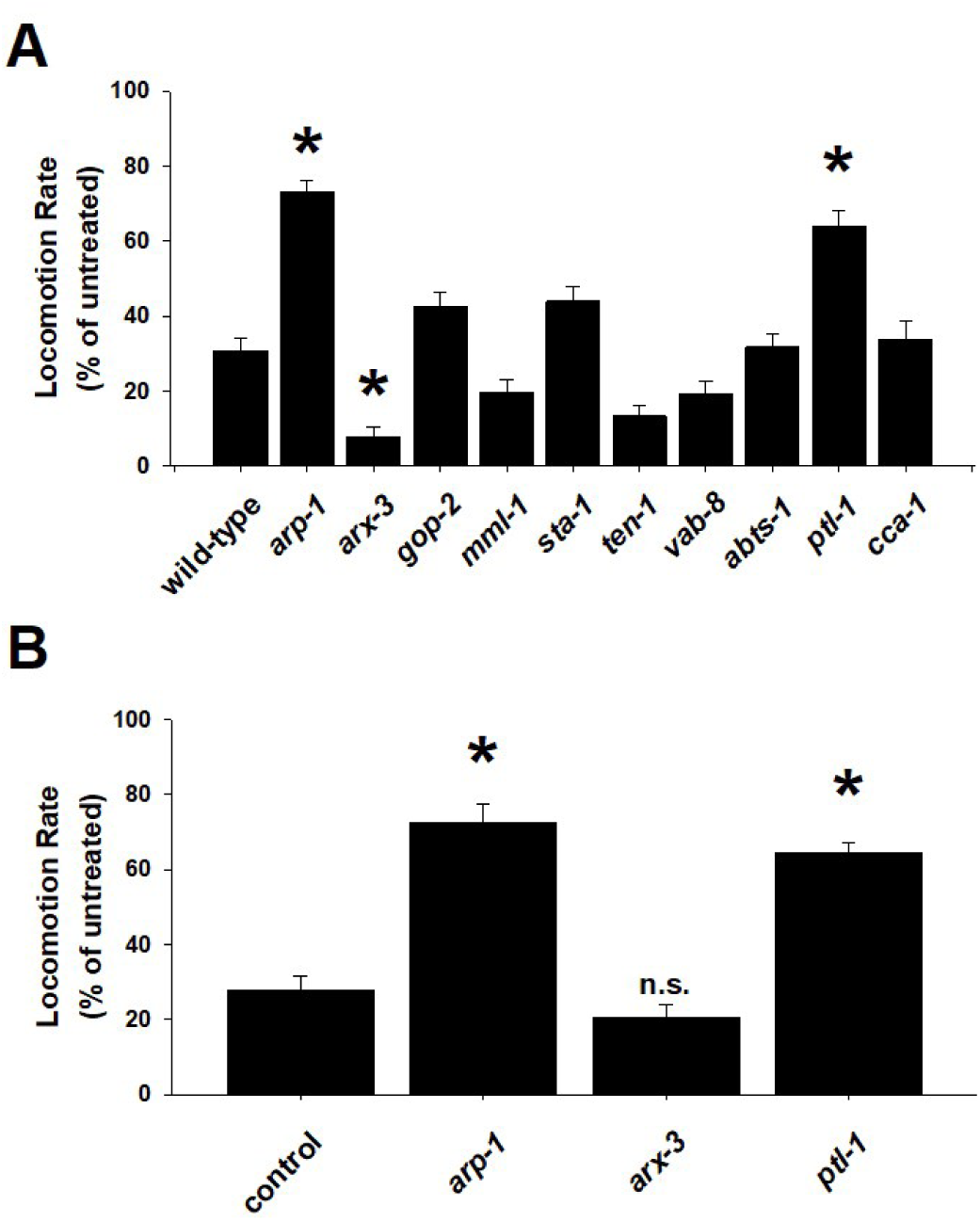
Quantification of alcohol phenotypes for *C. elegans* genes. (A) Nematodes with *loss-of-function* mutations in worm orthologues of *ACTR1B* (*arp-*1), *ARPC1B* (*arx-3*), *GPN1* (*gop-2*), *MLXIPL* (*mml-1*), *STAT6* (*sta-1*), *TENM2* (*ten-1*), *KIF26A* (*vab-8*), *SLC4A8* (*abts-1*), *MAPT* (*ptl-1*) and *SCN8A* (cca-1) were acutely exposed to ethanol and the resultant effect on rate of locomotion (thrashes per minute) was quantified. In comparison with Bristol N2 wild-type worms, significant differences were identified for *arp-1*, *arx-3* and *ptl-1*. Data is presented normalized to locomotion rate of untreated worms. *P<0.01. (B) RNAi confirmation of positively identified genes involved in alcohol phenotypes. In comparison to controls, RNAi knockdown of worm orthologues of *ACTR1B* (*arp-*1) and *MAPT* (*ptl-1*) phenocopied the *loss-of-function* mutations, whereas *ARPC1B* (*arx-3*) knockdown had no effect. *P<0.01. n.s., not significant.

### Secondary analysis

The Phewas analysis (**Table 3**) on the three genetic variants within *WDPCP*, *TENM2*, and *GPN1* showed a link between rs10078588 (*TENM2*) and food and liquid intake (beef, oily fish, fresh fruit, pork, bread, alcohol, water, coffee, salt) as well as trunk fat percent and smoking. rs13032049 (*WDPCP*) was linked to fresh fruit intake, salt and coffee intake, adiposity, angina pectoris, myocardial infarction, and depression. rs2178197 (*GPN1*) was linked to hypertension, hematologic traits, and body impedance. All three SNPs showed association with disorders of lipoprotein metabolism and other lipidaemias (ICD code E78) within the UK Biobank (**Table 4**). We finally performed a Mendelian randomization analysis on expression of *WDPCP* gene and liver conditions (**Table 5**) and identified a link between expression of this gene with liver fibrosis and cirrhosis (β=-0.20; 95% CI=-0.39, -0.01; *P*-value=0.04) and liver and bile duct cancer (β=0.0003; 95% CI=3.27×10^-05^, 5.85×10^-04^; *P*-value=0.02). We also observed a suggestive link with the changes in liver fatty acid-binding protein, and liver enzyme levels (**Table 5**).

**Table 3.**
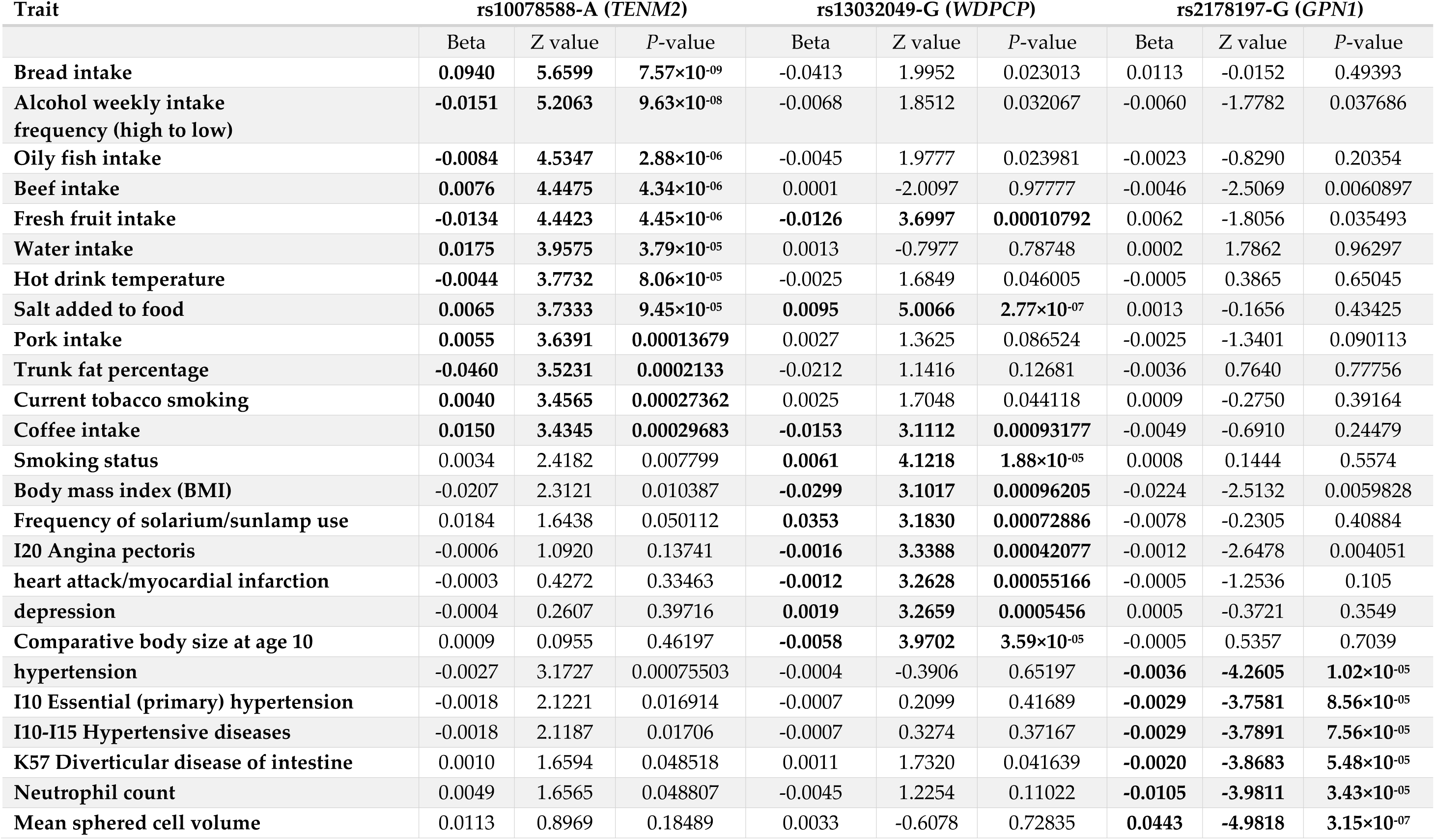
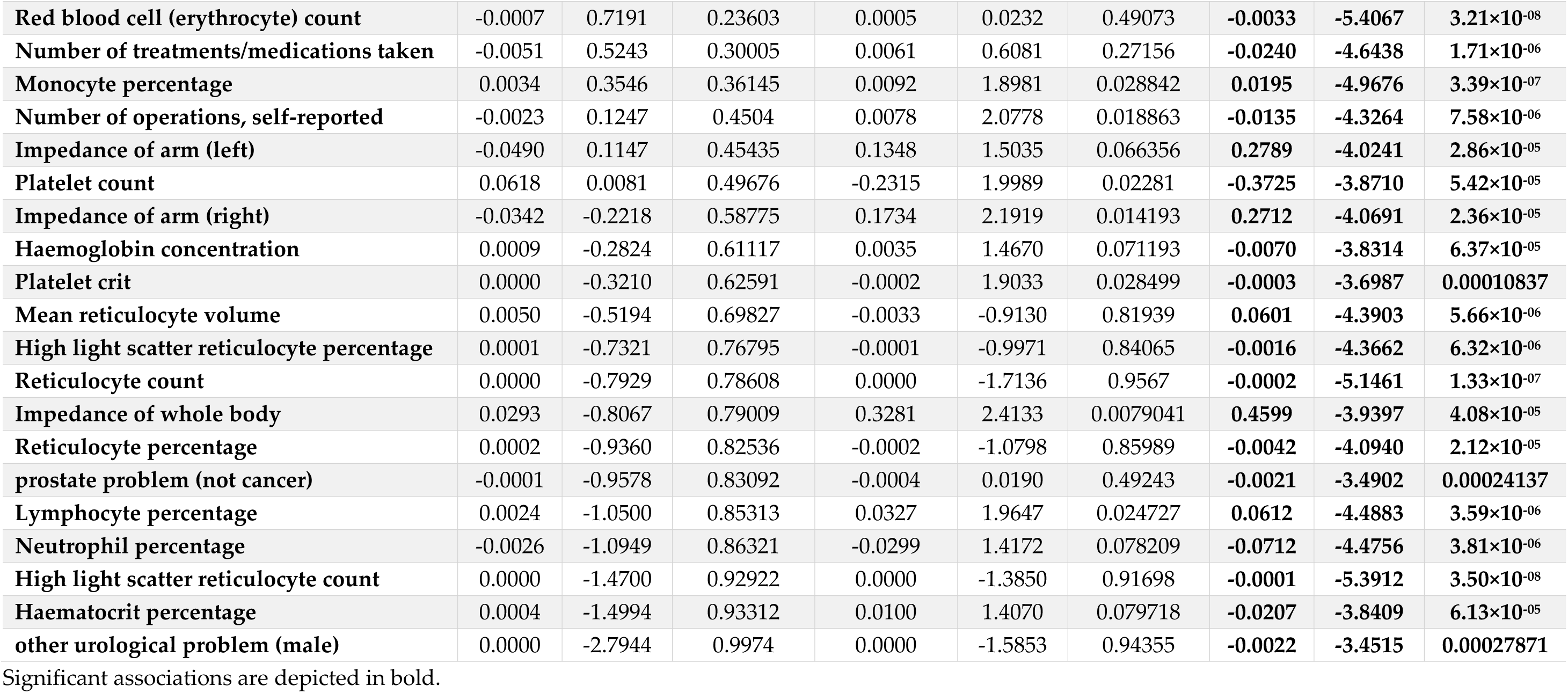
Overview of the significant associations between SNPs in *WDPCP*, *TENM2*, and *GPN1* with Phewas traits within the UK Biobank Edinburg Gene Atlas.

**Table 4.** Overview of the association of SNPs in WDPCP, TENM2, and GPN1 with disorders of lipoprotein metabolism and other lipidaemias (ICD code E78) within the UK Biobank Edinburgh Gene Atlas.

| SNP | chromosome | Base pair location | Effect allele | Beta (effect estimate) | 95% CI lower bound | 95% CI upper bound | P-value | MAF | HWE |
| --- | --- | --- | --- | --- | --- | --- | --- | --- | --- |
| **rs10078588** | 5 (*TENM2*) | 166816176 | A | -0.00134 | -2.49×10<sup>-04</sup> | -2.44×10<sup>-03</sup> | 0.016 | 0.474444 | 0.524 |
| **rs13032049** | 2 (*WDPCP*) | 63581507 | G | -0.00138 | -1.59<sup>-04</sup> | -2.60×10<sup>-03</sup> | 0.027 | 0.278928 | 0.5096 |
| **rs2178197** | 2 (*GPN1*) | 27860551 | G | -0.00144 | -3.36×10<sup>-04</sup> | -2.55×10<sup>-03</sup> | 0.011 | 0.424097 | 0.2078 |

Results obtained from the Edinburgh Gene Atlas. CI, Confidence Interval; SNP, Single Nucleotide Polymorphism; MAF, minor allele frequency; HWE, Hardy Weinberg Equilibrium.

**Table 5.** Overview of the significant results from inverse variance weighted Mendelian randomisation analysis for the effect of gene expression of ENSG00000115507 on liver traits using the MRC IEU OpenGWAS data infrastructure^37^.

| **Liver trait** | **Number of SNPs** | **Effect Estimate** | **Standard Error** | ***P* value** |
| --- | --- | --- | --- | --- |
| **Fatty acid-binding protein, liver** | 6 | -0.1053 | 0.0554 | 0.0576 |
| **Liver enzyme levels (alanine transaminase)** | 6 | -0.0017 | 0.0014 | 0.2355 |
| **Fibrosis and cirrhosis of liver** | 6 | -0.1972 | 0.0967 | 0.0415 |
| **Liver & bile duct cancer** | 6 | 0.0003 | 0.0001 | 0.0284 |

## Discussion

In this study we used data from humans, *C. elegans*, and *Drosophila* and identified a link between genes implicated in alcohol consumption and lipid metabolism. We identified that alcohol consumption changes the metabolites within linoleic acid (LNA) and alpha linolenic acid (ALA) metabolism pathway (LNA/ALA) and demonstrates causal effect on changes in several lipid metabolites. We highlighted that change of function of the genes implicated in alcohol consumption leads to changes in ethanol consumption, sedation after exposure to ethanol vapor, and changes in accumulation of fat in *Drosophila* as well as changes in locomotion rate after exposure to ethanol in *C. elegans.* Our results demonstrate that three alcohol implicated genes namely *WDPCP* (*frz*), *TENM2* (*Ten-m*), and *GPN1* (*CG3704*) might be involved in fat accumulation.

In our study, we used alcohol associated genetic variants to (1) explored the alcohol-induced biological changes in human metabolites and (2) explore alcohol-induced biological effect of genes annotated to alcohol associated genetic variants in *C. elegans* and *Drosophila melanogaster*. Of alcohol implicated genes that we investigated in *C. elegans*, *ACTR1B* (*arp-*1) and *MAPT* (*ptl-1*), show significant effects on the worms’ locomotion upon acute exposure to ethanol. In addition, *TENM2* (*Ten-m*) shows significant effects on ethanol consumption in *Drosophila* and apart from *MLXIPL* (*mondo*), RNAi lines for all the genes investigated in *Drosophila* change the time to sedation from ethanol.

The observed effect of *WDPCP* (*frz*) knockdown on the changed TAG levels occurred under exposure to ethanol and followed similar patterns as *hppy* mutants. This implies that minimizing alcohol consumption could reduce fat accumulation and thus reduce the risk of ALD. In our further MR analysis, we used publicly available databases derived from GWAS in human population and showed a link between the gene expression of *WDPCP* and liver fibrosis and liver cirrhosis. The analysis also showed a suggestive link with the liver fatty acid binding protein that is involved in the metabolism of lipids ^28^. This evidence suggests that *WDPCP* might be an important gene involved in the pathway between alcohol consumption, accumulation of fat and liver fibrosis. This could have public health implications in terms of identification of high-risk groups and targeting preventive measures as well as drug development. More studies in vivo and in vitro are needed to focus on *WDPCP* and provide more details on its role in lipid metabolism and liver pathologies.

Changes in TAG levels of *Drosophila* occurred with exposure to normal food rather than ethanol for RNAi knockdown of *TENM2* (*Ten-m*) and *GPN1* (*CG3704*) indicating that loss of function of these genes could have a direct role in accumulation of fat in liver independent of exposure to ethanol. RNAi knockdown of both genes shows increased tolerance to sedative effect of ethanol which could justify the effect of these genes on a more frequent alcohol consumption in human, possibly due to alcohol tolerance. *GPN1* is located on chromosome 2p23.3 and the encoded protein is implicated in regulation of TGFβ superfamily signaling ^29^ that is demonstrated to play a role in obesity ^30^, accumulation of fat in the liver ^31^ as well as regulation of *ADH1* gene that enhances alcohol-induced liver damage and lipid metabolism ^32^. The existing evidence alongside our findings on the role of *GPN1* in alcohol consumption and lipid metabolism in *Drosophila* implies that *GPN1* might play a role upstream of TGFβ in the regulation of metabolism of alcohol and lipids. Further studies are needed to highlight the relationship between *GPN1* and TGFβ in alcohol consumption and alcohol-induced liver damage.

*TENM2* (*Ten-m*) is located on chromosome 5q34, and the encoded protein is involved in cell adhesion ^33^. *TENM2* is found to be highly enriched in white adipocyte progenitor cells ^34^. *TENM2* deficiency in human fat cells leads to expression of UCP1, the primary marker of brown adipose tissue ^35^. Genetic variants in *TENM2* have shown to be linked to obesity ^36^. Our secondary analyses confirmed an association between the genetic variant in *TENM2* and excess of food and liquid intake which suggests the link between *TENM2* and alcohol consumption could also be due to systematic increase in consumption of all food and beverages rather than alcohol alone. The evidence in this study alongside the existing literature highlights that the observed effect of *TENM2* (*Ten-m*) RNAi knockdown on the changes in TAG levels in *Drosophila* could potentially be related to biological pathways implicated in adipose tissue rather than pure liver-related pathways.

One strength of our study is in that we performed our analyses in human and two different model organisms that allows for a more comprehensive insight into biological mechanisms involved in the function of alcohol consumption genes under different biological scenarios. A second strength of this study is in the use of RNAi technique which provides insight into the function of genes and what biological manifestation they would have when exposed to ethanol. The third strength of our study is in the use of CAFE assay that allows for investigation of food and alcohol consumption in *Drosophila* in a more controlled environment. In our CAFE assay, each experimental box per genotype contained both normal food and ethanol food (food supplemented with 15% ethanol) providing the insects with a choice. Another strength of our study is in that in our TAG levels experiments, we made our conclusions based on the comparisons between flies (RNAi vs. control) that were exposed to identical food and environmental conditions which increases the robustness of our conclusions. Finally, we combined the results with human studies to get better insight into the link between alcohol consumption and lipid metabolites.

Although the genetic variants used for our investigations were originally found in human studies and we also performed a metabolomics analysis between alcohol consumption and circulating metabolites in human population, the main conclusions are made based on the effect of the genes in model organisms and the results of this study might not directly generalisable to patients and the public without performing further population studies.

## Conclusion

We identified that alcohol associated genes may be involved in metabolism of lipids and accumulation of fat in liver. Our study highlights three genes *WDPCP, TENM2,* and *GPN1* that may be involved in accumulation of fat (in liver or adipose tissue). Of these genes *WDPCP* exhibits its effects on fat accumulation in *Drosophila* with exposure to ethanol. The gene expression of *WDPCP* in human population supports a link to liver fibrosis. Further studies are necessary to investigate the role of this gene in ALD.

## Acknowledgements and sources of funding

R.P. was supported by Rutherford Fund fellowship from the Medical Research Council (MR/R0265051/1 and MR/R0265051/2). B.A., X.J., and F.O. were supported by Rutherford Fund from Medical Research Council MR/R0265051/2. R.M. was funded by the President’s PhD Scholarship from Imperial College London. PE is Director of the MRC Centre for Environment and Health and acknowledges support from the Medical Research Council (MR/S019669/1). PE also acknowledges support from the UK Dementia Research Institute, Imperial College London (UKDRI-5001), Health Data Research UK London (HDRUK-1004231) and the British Heart Foundation Imperial College London Centre for Research Excellence (BHF-RE/18/4/34215). The Airwave Health Monitoring Study was funded by the UK Home Office (780-TETRA, 2003-2018) and is currently funded by the MRC and ESRC (MR/R023484/1) with additional support from the NIHR Imperial College Biomedical Research Centre in collaboration with Imperial College NHS Healthcare Trust. R.C.P is supported by the UK Dementia Research Institute (UKDRI-5001), which receives its funding from UK DRI Ltd, funded by the UK Medical Research Council, Alzheimer’s Society and Alzheimer’s Research UK. Work in LMM’s laboratory is supported by the UK Medical Research Council, intramural project MC_UU_00025/3 (RG94521). The views expressed are those of the authors and not necessarily those of the sponsors. We thank Prof. Ulrike Heberlein, (Janelia Research Campus, Virginia, USA) for generously providing us the hppy17-51 fly lines.

---

**Supplementary Figure S1:**
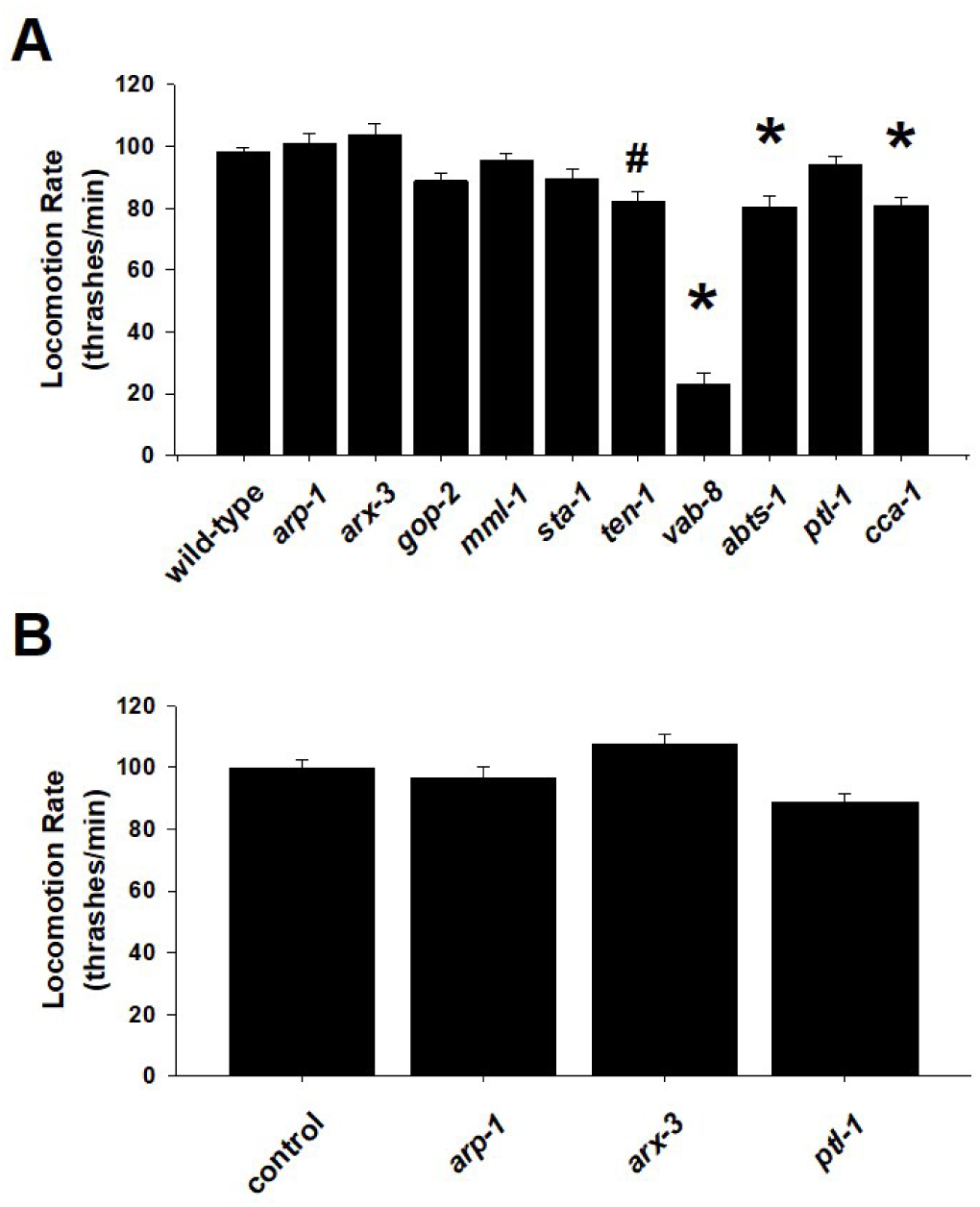
Basal locomotion rates. (A) Nematodes with *loss-of-function* mutations in worm orthologues of *ACTR1B* (*arp-*1), *ARPC1B* (*arx-3*), *GPN1* (*gop-2*), *MLXIPL* (*mml-1*), *STAT6* (*sta-1*), *TENM2* (*ten-1*), *KIF26A* (*vab-8*), *SLC4A8* (*abts-1*), *MAPT* (*ptl-1*) and *SCN8A* (cca-1) were quantified for locomotion rate (thrashes per minute). In comparison with Bristol N2 wild-type worms, significant differences were identified for *ten-1*, *vab-8*, *abts-1* and *cca-1*. *P<0.01. #P<0.05. (B) Quantification of locomotion rate for worms subjected to RNAi knockdown. In comparison to controls, RNAi knockdown of *arp-1*, *arx-3* or *ptl-1* had no effect on basal locomotion rate.

