## Supplemental Information for "Evidence for Involvement of *WDPCP* Gene in Alcohol Consumption, Lipid Metabolism, and Liver Cirrhosis"


**Supplementary Methods**

*Study Population.* The Airwave Health Monitoring Study (the Airwave Study) was first established in 2004 as a large-scale occupational cohort of police officers. A total of 53,114 participants were enrolled by the end of baseline recruitment in March 2015. Initially, it aimed to investigate the health outcomes related to the use of Terrestrial Trunked Radio (TETRA).

This cohort has also been expanded to investigate the health of workforces in more general. The rationale, design, and methods of this study can be found elsewhere [1]. The Airwave Health Monitoring Study was approved by the National Health Service Multi-site Research Ethics Committee (MREC/13/NW/0588).

*Metabolomic assays.* A random sample of 2063 participants in the Airwave study were assayed for various metabolites. Blood samples were collected at the screening visit (with informed consent) and stored at -80 °C, before being transferred to a biorepository facility and stored in vapor phase liquid nitrogen. The metabolomic assays were performed using Liquid chromatography-mass spectrometry (LC-MS) at the Imperial Phenome Centre (London, UK). Lithium heparin was utilized as an anticoagulant for the LC-MS plasma samples. Metabolomics data was acquired using Ultra Performance Liquid Chromatography-Mass Spectrometry (UPLC-MS) and Quality Control (QC) methods that were previously described [2-4]. This involved preparing a pooled study reference sample for each population and regular analysis of QC samples throughout data acquisition. Additionally, mixtures of authentic reference standards were added to the study reference, long-term reference, and study samples used in the UPLC-MS analysis to enable targeted monitoring of data quality during acquisition. Plasma samples were prepared and underwent UPLC-MS profiling analysis for lipids and small metabolites as previously described by Izzi-Engbeaya and colleagues [3]. The only deviation from the methodology in Izzi-Engbeaya and colleagues was that 100µl of samples was used without dilution prior to addition of isopropanol for the lipidomics analyses. Batches of 80 samples were prepared into 96-well plates. Each sample was mixed with four parts of 4°C isopropanol, incubated at 4°C, centrifuged, and the supernatant aliquoted into a 96-well plate. All analyses were acquired on Acquity UPLC systems coupled to Xevo G2-S ToF mass spectrometers (Waters Corporation, Milford, Massachusetts, United States). Three platforms were used for these samples: reverse phase in positive ionization mode which detects largely lipid species (Lipid Positive mode, LPOS), reverse phase in negative ionization mode which detects largely lipid species (Lipid Negative mode - LNEG) and Hydrophilic Interaction Chromatography (HILIC) in positive mode which detects largely small and polar metabolites (HILIC Positive mode, HPOS).

*Metabolomics Data processing.* Peak picking was performed using Bioconductor R-package XCMS [5]. Drift correction was achieved using a previously described method [6]. Then, we replaced negative values with zeros, and added one before log-transforming the data. Metabolomic features were filtered based on retention time to exclude non-retained and cleaning phase features - only features in the following retention times (in minutes) were accepted: HPOS (0.5-7); LNEG (0.3-9.5); LPOS (0.45-12). Principal components analysis (PCA) was used to identify outlier samples, which were then excluded. Furthermore, we excluded single-point values that were more than 5 Median Absolute Deviations (MAD) from the median. We used 10 principal components from genome-wide scans to adjust for population stratification. Finally, we transformed the data into z-scores using median and MAD so that we had comparable intensities across studies.

*UPLC-MS metabolite annotation.* Lipid annotation was initially completed by matching accurate mass fragmentation measurements to reference spectra from online databases (LIPID

MAPS, Metlin, HMDB) and previous publications. Where chemical reference materials were commercially available (Avanti Polar Lipids, Sigma Aldrich, Cayman Scientific), they were used to generate definitive molecular identification by direct matching of chromatographic and spectral qualities (including accurate mass, MS/MS spectra, and isotopic distribution) to those observed in the profiling data.

*Genotyping and imputation.* In the Airwave study [1], genotyping was conducted using Illumina Infinium HumanExome-12v1-1 BeadChip Array. QC steps were performed to remove any samples with high missingness (>3%) or outlier heterozygosity rates (>3SD from the batch mean), duplicates (ID-based or genotype-based) or presented with high degree of relatedness. Markers with genotype call rate <98%, significant deviation from Hardy-Weinberg equilibrium ($P < 1 \times 10^{-5}$), and low MAF (<1%) were removed. Genotype data was imputed with 1000 Genomes phase 3 reference panel.

*GWAS on metabolomics.* We performed genome-wide association studies (GWAS) in the Airwave Study to obtain genetic association estimates for alcohol-related SNPs and selected metabolomic features. Untargeted Mass-Spectrometry (MS) was performed in heparin plasma samples of the Airwave Study, producing three separate datasets, namely hydrophilic interaction liquid chromatography (HILIC) positive (HPOS), lipid positive (LPOS), and lipid negative (LNEG). For each metabolomic dataset, we used principal component analysis (PCA) to identify outlier samples and exclude them. The data was residualised using first ten principal components to account for population stratification. We transformed the data into z-scores using median and median absolute deviations (MAD). Values that were more than 5 MAD from the median were removed and imputed using k-nearest neighbors' imputation [7]. GWAS were conducted for each metabolomic feature with adjustment for age and sex (N=1,970). Due to the high number of metabolomic features, GWAS were performed using the high-dimensional association analyses (HASE) framework, which applies matrix operation and removes redundant calculation in high-dimensional association analyses to improve computational efficiency [8].
